# Detection of statistically significant network changes in complex biological networks

**DOI:** 10.1101/061515

**Authors:** Raghvendra Mall, Luigi Cerulo, Halima Bensmail, Antonio Iavarone, Michele Ceccarelli

## Abstract

**Motivation:** Biological networks contribute effectively to unveil the complex structure of molecular interactions and to discover driver genes especially in cancer context. It can happen that due to gene mutations, as for example when cancer progresses, the gene expression network undergoes some amount of localised re-wiring. The ability to detect statistical relevant changes in the interaction patterns induced by the progression of the disease can lead to discovery of novel relevant signatures.

**Results:** Several procedures have been recently proposed to detect sub-network differences in pairwise labeled weighted networks. In this paper, we propose an improvement over the state-of-the-art based on the Generalized Hamming Distance adopted for evaluating the topological difference between two networks and estimating its statistical significance. The proposed procedure exploits a more effective model selection criteria to generate p-values for statistical significance and is more efficient in terms of computational time and prediction accuracy than literature methods. Moreover, the structure of the proposed algorithm allows for a faster parallelized implementation. In the case of dense random geometric networks the proposed approach is 10−15x faster and achieves 5-10% higher AUC, Precision/Recall, and Kappa value than the state-of-the-art. We also report the application of the method to dissect the difference between the regulatory networks of IDH-mutant versus IDH-wild-type glioma cancer. In such a case our method is able to identify some recently reported master regulators as well as novel important candidates.

**Availability:** The scripts implementing the proposed algorithms are available in R at https://sites.google.com/site/raghvendramallmlresearcher/codes.

## 5 Introduction

The omnipresence of complex networks is reflected in wide variety of domains including social networks (Jin *et al.*, 2013;Mislove *et al.*, 2007), web graphs (Broder *et al.*, 2000), road graphs (Erath *et al.*, 2009), communication networks (Kesidis, 2007), financial networks (Boginski *et al.*, 2005) and biological networks (Ideker *et al.*, 2002;Keller *et al.*, 2009;Nacu *et al.*, 2007). Although we focus on biological networks many aspects of the method proposed in this paper can also be applied for networks in other contexts. In cancer research comparisons between gene regulatory networks, protein interaction networks, and DNA methylation networks is performed to detect difference between conditions, such as healthy and disease (Dehmer and Emmert-Streib, 2008;*D*’haeseleer *et al.*, 2000). This can lead to discovery of biological pathways related to the disease under consideration, and, in case of cancer, the gene regulatory changes as the disease progresses (Wallace *et al.*, 2011;Ahern *et al.*, 2016).

A central problem in cell biology is to model functional networks underlying interactions between molecular entities from high throughput data. One of the main questions is how the cell globally changes its behavior in response to external stimuli or as the effect of alterations such as driver somatic mutations or changes in copy number. Signatures of differentially expressed and/or methylated genes are the downstream effect of the de-regulation of the global behavior of the cell in different conditions such as cancer subtypes. Therefore, it is argued that driver mutations activate functional pathways described by different global re-wiring of the underlying gene regulatory network.

The identification of significant changes induced by the presence or progression of the disease can help to discover novel molecular diagnostics and prognostic signatures. For example, we have recently shown in (Ceccarelli *et al.*, 2016) that the majority of malignant brain tumors can be divided two main macrocategories according to the mutation of the gene IDH which can be further divided in seven molecular and clinically distinct groups. These two macrogroups are characterized by highly different global expression and epigenomic profiles. Hence, one of the main questions to understand the molecular basis of diseases is how to identify significant changes in the regulatory structure in different conditions, in a similar way we analyze differentially expressed genes in different conditions.

Various techniques have been developed to compare two graphs including graph matching and graph similarity algorithms (Brandes and Eriebach, 2005;Lena *et al.*, 2013;Yang and Sze, 2007). However, the problem addressed in this paper is different from popular graph theory problems including graph isomorphism (Ramana *et al.*, 1994) and sub-graph matching (Shervashidze *et al.*, 2011). Here the goal is to identify statistically significant differences between two weighted networks (with or without labels) under the null hypothesis that the two networks are independent.

One common statistic used to distinguish one graph from another is the Mean Absolute Difference (MAD) metric defined as: 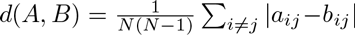 where *a*_*ij*_ and *b*_*ij*_ are edge weights corresponding to the topology of networks *A* and *B*. This distance measure is equivalent to the Hamming distance (Hamming, 1980) and has been extensively used in literature to compare networks (Butts and Carley, 1998; Gill *et al.*, 2010). Another statistic used to test association between networks is the Quadratic Assignment Procedure (QAP) defined as: 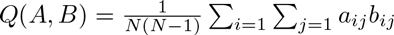. The QAP metric (Mantel, 1967;Hubert, 1987) is used in a permutation-based procedure to differentiate two networks. Ruan *et al.* (2015) showed that these metrics are not always sensitive to subtle topological variations.

Our aim is to detect statistically significant differences between two networks under the premise that any true topological difference between the two networks would involve only a small set of edges when compared to all the edges in the network. Recently, a Generalized Hamming Distance (GHD) based method was introduced to measure the distance between two labeled graphs (Ruan *et al.*, 2015). It was shown in Ruan *et al.*, (2015) that the GHD statistic is more robust than MAD and QAP metrics for identifying subtle variations in the topology of paired networks.

The authors in Ruan *et al.*, (2015) propose a non-parametric test for network comparison where they provide conditions for asymptotic normality such that p-values can be obtained in closed-form. They also propose a differential sub-network identification technique namely dGHD. The advantage of this technique is that it provides closed-form solution for p-values for the sub-network left after iterative removal of the least differential nodes unlike previous differential network analysis techniques (Gill *et al.*, 2010;Fuller *et al.*, 2007;Ha *et al.*, 2015). We propose an extension of dGHD, namely Closed-Form approach that, exploiting the conditions for asymptotic normality (Ruan *et al.*, 2015), is computationally cheaper and attains better prediction performance than the Original (dGHD) algorithm. Computational efficiency and prediction accuracy is crucial in cancer contexts where networks have a large number of nodes and the topological difference is associated to few driver genes.

The paper is organized as follows: Section 2 introduces the improved algorithm to detect statistically significant sub-network differences; Section 7 defines the experimental procedures adopted to evaluate the proposed method and discusses the results of the experiments; Section 8 reports the results of the application of the proposed procedure in the context of glioma cancer showing that we identify some of the relevant driver genes known in literature; and Section 9 concludes the paper drawing future directions.

## 6 Methods

### 6.1 Preliminaries on Generalized Hamming Distance

The Generalized Hamming Distance is a way to estimate the distance between two graphs (Ruan *et al.*, 2015). Let *A* = (*V*, *E*_*A*_) and *B* = (*V*, *E*_*B*_) two graphs, with the same set of nodes *V* = {1,…*N*}, and different sets of edges, *E*_*x*_ representing the set of edges in the network *X*. The Generalized Hamming Distance (GHD) is defined as:

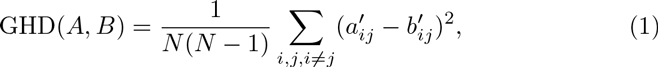

where *a*′_*ij*_ and *b*′_*ij*_ are mean centered edge-weights defined as:

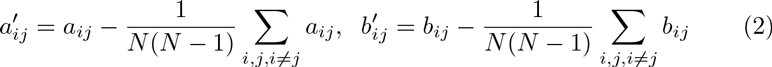

The edge weights, *a*_*ij*_ and *b*_*ij*_, depend on the topology of the network and provide a measure of connectivity between every pair of nodes *i* and *j* in *A* and *B*. Different metrics have been adopted to measure the connectivity between pair of nodes including topological overlap (TO) measure (Zhang and Horvath, 2005;Allen *et al.*, 2012), cosine similarity and pearson correlation (Deshpande *et al.*, 2013). In our experiments, we used the cosine similarity metric to create the topological network corresponding to graph *A* and *B*. We utilized the cosine similarity metric to capture first order interactions between the nodes in the network. This is due to its ease of implementation for large scale sparse networks using set operations. The cosine metric has nearly perfect correlation with TO measure (Supplimentary Fig 1). Hence it can be used as a replacement to TO measure, adopted in Ruan *et al.*, 2015), while constructing the topological networks for graphs *A* and *B*.

The problem of detection differential sub-networks is posed as an inferential problem with a statistical hypotesis test under the null hypothesis 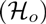: *Graphs A and B are independent*. The null distribution can be obtained with a permutation test, as shown in (Ruan *et al.*, 2015), by constructing a sampling distribution of GHD computed between *A*_*π*_ = (*V*_*π*_, *E*_*A*_) and B = (*V*, *E*_*B*_), where *V*_*π*_ is a permuted version of the set of vertex *V*. By keeping *B* as reference network, each permutation consists of shuffling the labels of the nodes in *A* while keeping the edges unchanged.

The authors in Ruan *et al.*, (2015) demonstrated that GHD computed on the permuted version follows a normal distribution. So, by providing conditions for asymptotic normality one can efficiently calculate the p-value circumventing the computationally expensive of an empirical permutation test. This can be shown as:

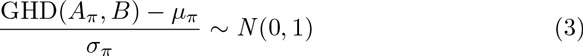

Here *μ_π_* is the asymptotic value of the mean GHD and *σ*_*π*_ is the asymptotic value of the standard deviation of GHD computed between *A*_*π*_ and *B*. In order to calculate the *μ_π_* and *σ*_*π*_ values we define:

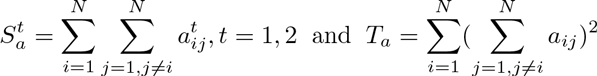

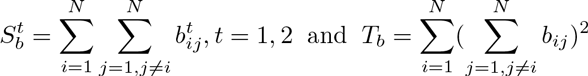

Here 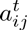 and 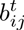 are the edge weights with the power *t*. Furthermore, we require the following terms:

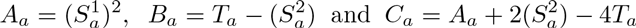

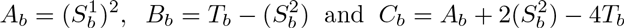

Using these definitions the closed-form expression for mean *μ_π_* and variance 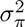 are expressed as:

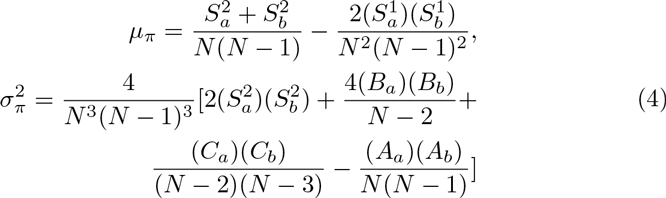

Given a significance threshold *α* (e.g. 0.01), p-values > *α* indicate that there is no sufficient evidence to reject the null hypothesis 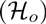 that graphs *A* and *B* are independent. Hence, higher p-values indicate more probability that the two graphs under consideration are independent.

### 6.2 Differential sub-network detection with GHD

The GHD distance is able to tell us to what extent are two graphs different but is not able to identify which parts of the graph are similar and which are different. In this work, we are interested in detecting which part of the graphs contribute to make the two graphs different. We call such different sub-graphs *differential sub-networks*.

The notion of differential sub-networks is based on the idea that when comparing two networks only a subset of edges would present altered interaction. The goal is to identify the set of nodes, namely *V*\*, associated with such a subset of edges and the p-values *p*\* corresponding to the nodes in *V*\*. This goal, formulated as a statistical test, requires that for such a subset *V*\* there is no sufficient evidence to reject the null hypothesis that the corresponding sub-networks *A*\*(*V*\*,*E*_A*_) and *B*\*(*V*\*,*E*_B*_) are statistically independent.

The idea here is to adopt an iterative technique to identify the set of nodes *V*\* that contributes more to the difference. We start from the dGHD algorithm proposed in Ruan *et al.*, (2015). The algorithm measures the edge connectivity with topological overlap metric and benefits from the closed-form solution of p-value (Equations 4). In the dGHD algorithm, an iterative procedure is followed where during each iteration the change in centralized GHD (cGHD) i.e. cGHD = GHD(A,B) −*μ_π_* is estimated after the removal of one node. The node corresponding to which the change in cGHD value (i.e. difference in cGHD value before and after removal of a node) is maximum is removed. The GHD statistic is computed for remaining sub-networks and the p-value is estimated. This process is repeated till a user specified minimal set size is reached or it is no-longer possible to have closed-form representation for p-values which happens for N ≤ 3 as shown in equation 4. The p-values are adjusted for multiple testing by controlling the false discovery rate (Benjamini and Yekutieli, 2001).

The dGHD algorithm suffers from the following limitations: a) During the *i*^*th*^ iteration, the GHD measure is calculated N − *i* times on different sub-graphs with an overall time complexity 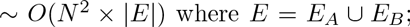 b) The algorithm is prone to discovery of more false positives since it uses the change in cGHD as a model selection criterion. We overcome such limitations by proposing the following improvements:

1. *Remove nodes by exploiting the Closed-Form.* We use the idea that nodes which have similar topology in networks *A* and *B* will contribute the least to cGHD. So, we first calculate the closed-form contribution of each node in cGHD once using equation 5 and then iteratively remove nodes with least contributions. However, this process is continued till we observe that the p-value of the remaining sub-network becomes greater than a threshold *θ*.
2. *Using a different model selection criterion*. Once the p-value reaches *θ*, we follow a procedure similar to the dGHD algorithm but use the more intuitive criterion of selecting the node that when removed makes the cGHD value maximum rather than using the change in the cGHD value (before and after removal of a node) as a model selection criterion. By using this model selection criterion, we iteratively identify and remove that node whose contribution is least in the cGHD. The advantage of the Closed-Form approach is that we significantly reduce the computational complexity and improve the predictive performance. A simple alternative to the Closed-Form approach would be to sort all the nodes based on their contribution to cGHD and thus rank all the nodes based on their capability to differentiate the two networks with complexity (*O*(*N* log *N*)). However, then we will not be able to identify statistically different sub-networks between the two graphs as indicated in Ruan *et al.*, 2015).

#### 6.2.1 Closed-Form Approach

We propose a fast approach to perform differential sub-network analysis taking into consideration the contribution of each node in the GHD and *μ_π_*. Using equations 1 and 4 this can mathematically be represented as:

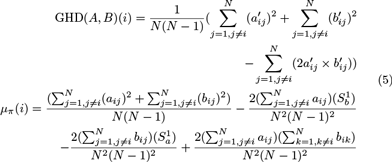

We observe that if we add the GHD(*A*, *B*)(*i*) and *μ_π_* (*i*) ∀*i*, we obtain GHD(*A*, *B* and *μ_π_*. We use the idea that nodes which have similar topology in networks *A* and *B* will contribute the least to centralized GHD, i.e. GHD(*A, B*) − *μ_π_*. We calculate the Closed-Form contribution of each node in the centralized GHD (cGHD) once using equation 5 and then iteratively remove nodes with least contribution to the cGHD, i.e. nodes having similar topology in graphs *A* and *B*. Thus, we calculate cGHD once and sort all the nodes based on their contribution to the cGHD metric.

This process is continued till we observe that the p-value of the remaining sub-network becomes greater than a threshold *θ*. Once the p-value reaches *θ*, we estimate 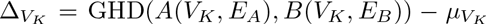 where 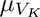 is the mean of the permutation distribution for the nodes (*V*_*K*_) of the remaining subnetwork. Furthermore, we define 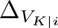 as the value of cGHD after removal of node *i*. We adopt a different model selection criterion than that proposed in Ruan *et al.*, (2015) to remove non-differential nodes. We use the intuitive criterion of selecting that node after removal of which the cGHD value becomes maximum, i.e. the node whose contribution was least significant in cGHD or the node which was most similar in terms of topology for the paired-graphs. Finally, the obtained p-values are adjusted for multiple testing by controlling the false discovery rate (Ruan *et al.*, (2015)Benjamini and Yekutieli, 2001). Provided the paired-graphs *A* and *B*, the calculation of 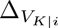 can be done independently for each *i*. Details of the Closed-Form method is provided in Algorithm 1. Table 1 summarizes the improvements with respect to the dGHD algorithm in terms of time complexity.

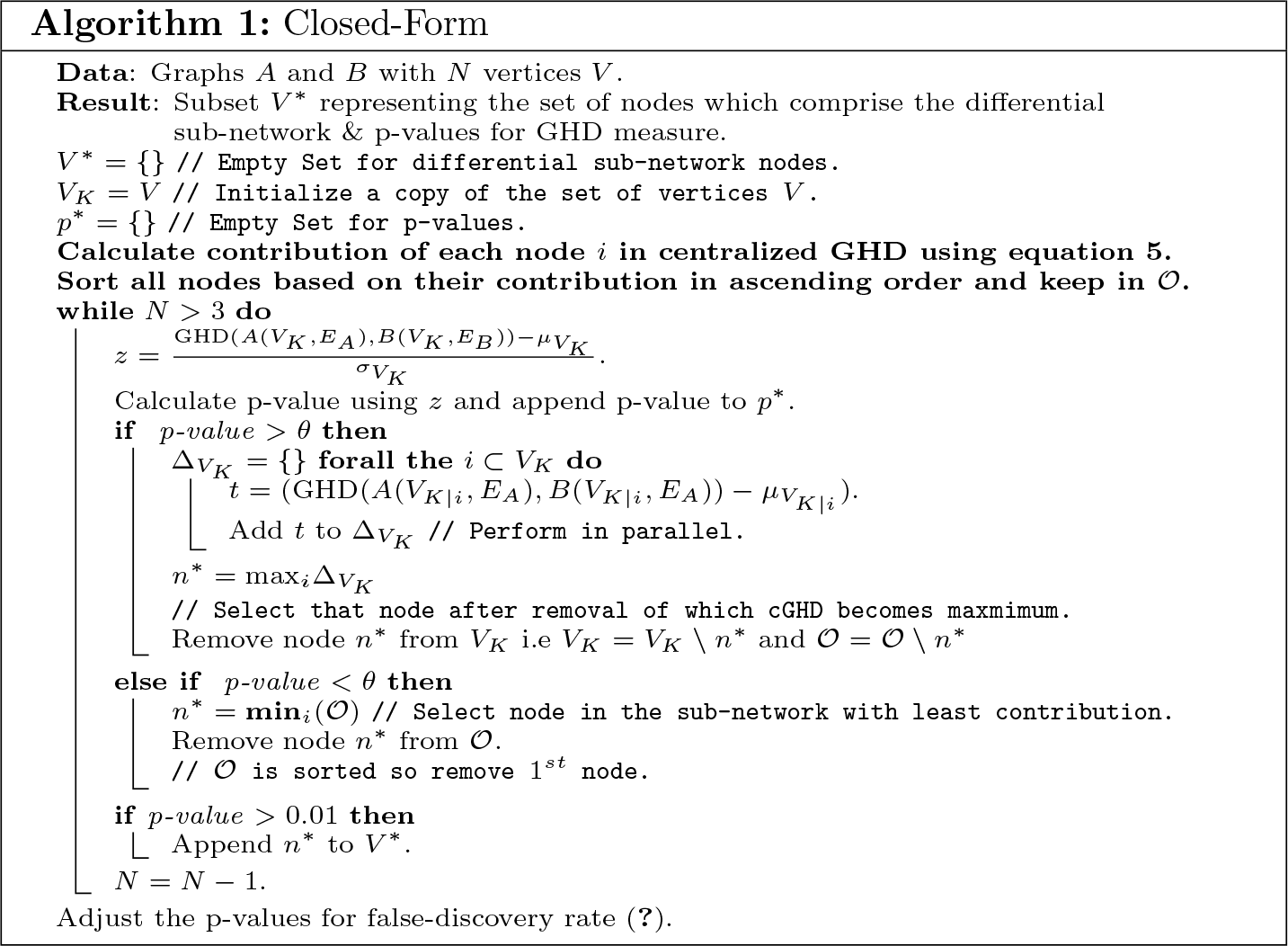

**Table 1:** Comparison of time complexity. Here *K* represents the number of nodes for which p-value is greater than *θ* and generally *K* ≪ *N*. An important remark is that the cGHD calculation after removal of each node can be done independently in parallel. So, in case we have *T* processors, the complexity of the proposed approach can be reduced ≈ linearly w.r.t. *T*.

| Computational Complexity |  |
| --- | --- |
| dGHD | Closed-Form |
| $O(N^2\vert E\vert )$ | $O(N\vert E\vert + N \log(N) + K^2\vert E\vert )$ |

#### 6.2.2 Alternative Procedure (Fast Approximation)

We propose an alternative procedure to the Closed-Form approach namely the Fast Approximation method where we first calculate the cGHD value without including the *i*^*th*^ node, ∀_*i*_ ∈ *V* once. This helps to estimate the cGHD value after removal of the *i*^*th*^ node and can be performed in parallel. Our aim is to quickly discard those nodes after removal of which the cGHD value becomes large thereby removing nodes which were contributing least to the cGHD value. This helps to reduce the dependence between the two sub-networks by removing nodes which have similar topology in graphs *A* and *B*. Again, the idea is motivated by the premise that only a subset of nodes will form the differential sub-networks in graph *A* and *B*.

In this approach, we iteratively discard those nodes after removal of which the cGHD value becomes maximal till the p-value for the remaining sub-network reaches a threshold *θ*. Once the p-value reaches *θ*, we return back to the procedure of estimating 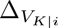 ∀_*i*_ ∈ *V*_*K*_ as described in the Closed-Form approach. We use the same model selection criterion of selecting that node after removal of which the cGHD value becomes maximum as used in the Closed-Form approach. We then adjust the obtained p-values for multiple testing by controlling the false discovery rate (Benjamini and Yekutieli, 2001). We refer to this technique as a Fast Approximation to the Original technique (dGHD (Ruan *et al.*, 2015)). We explain the Fast Approximation technique in detail in (Supplementary Algorithm 1).

From our experiments, we observe that the results of the Closed-Form approach and the Fast Approximation technique are identical. Although, in the case of Closed-Form approach, we calculate closed-form contribution of each node in the cGHD value and remove the node with least contribution, while in case of Fast Approximation we select that node after removal of which cGHD value becomes maximum, the ordered list O obtained for both the methods is identical. Moreover, the computational complexity of the Fast-Approximation technique is the same as that of Closed-Form approach.

## 7 Experimental Results

For all our experiments, we used the Closed-Form approach (since results obtained from Closed-Form and Fast-Approximation techniques are identical) and compare it with the dGHD method Ruan *et al.*, (2015).

### 7.1 Sensitivity to *θ*

In this experiment, we check the sensitivity of the proposed Closed-Form approach w.r.t. the heuristic *θ*. For this experiment, we first generated 100 random geometric (RG) networks. In a RG network nodes are generated by uniformly sampling *N* points on [0,1]^2^. An edge is then drawn between these points if the euclidean distance between the points is less than a parameter *d*. This parameter d controls the density of the RG network where smaller values of *d* result in sparse networks while larger values of *d* generates dense networks. In our case, we conducted experiments using two different settings. In the first setting, we use *d* = 0.15 while in the second case we use *d* = 0.3. For both the experiments we fix *N* = 250. For each value of d and for each generated RG network *A*, we permute the first 50 rows and columns of the network to generate network *B*. Therefore, the first 50 nodes in networks *A* and *B* form the true positives (TP).

In order to test the sensitivity of the proposed approach w.r.t. *θ*, we estimate the fraction of permuted nodes (TP) correctly identified by the Closed-Form method for various values of *θ*. We used a grid of *θ* values varying from Θ = {*e*^−50^,…,*e*^−250^} in multiplicative steps of *e*^−20^. The goal of this experiment is to show that the fraction of TPs identified w.r.t. various *θ* ∈ Θ remains nearly constant for smaller values of *θ*.

**Figure 1:**
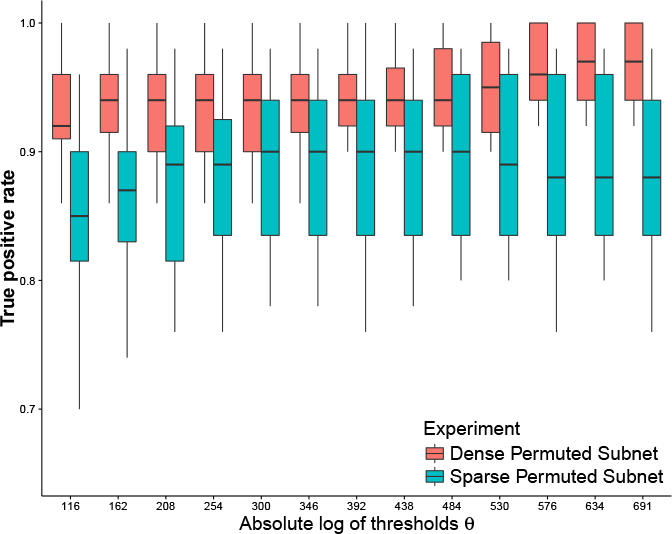
The boxplots represents the distribution of True Positives (TP) identified by Closed-Form approach for 100 random runs of the experiment.

Figure 1 shows the result for RG networks with density parameter *d* = 0.15 and *d* = 0.3. From Figure 1, we observe that the median fraction of permuted nodes identified by the proposed approaches increases slowly before it converges to a nearly constant value as we decrease the threshold 0 (i.e. increase absolute log of threshold *θ*). From this experiment, we conclude that:

> *The fraction of truly differential nodes (TP) identified by the proposed methods increases as we decrease the threshold 0 before it starts to converge for smaller values of threshold *θ**

We used the threshold *θ* = *e*^−250^ as heuristic for p-value cutoff in future experiments.

### 7.2 Predictive performance validation

The next simulation study that we carried out was to compare the predictive performance of the proposed approach w.r.t. the dGHD (Ruan *et al.*, 2015) technique. For this experiment, we generate 100 RG networks with *N* = 1,000. For the first experiment we fix the density parameter *d* = 0.3 and permute first 100 nodes in network *A* to obtain network *B*. Thus, these first 100 nodes form the differential sub-network for the paired networks *A* and *B*.

In the second case, we use the same density parameter *d* = 0.3 to generate the edges for network *A*. We then generate a small RG network with 100 nodes using density parameter *d*′ = 0.5. This small dense sub-network is then used to replace the network formed by first 100 nodes in the original network *A* to form network *B*. Thus, in the second experiment, these 100 nodes form the differential sub-network for the paired networks *A* and *B*. This kind of mechanism can appear in real-life networks, for example, in case of cancer the transcription activity of some set of genes might get enhanced or suppressed in patients resulting in more or fewer edges in a sub-network of the gene or DNA methylation network.

**Figure 2:**
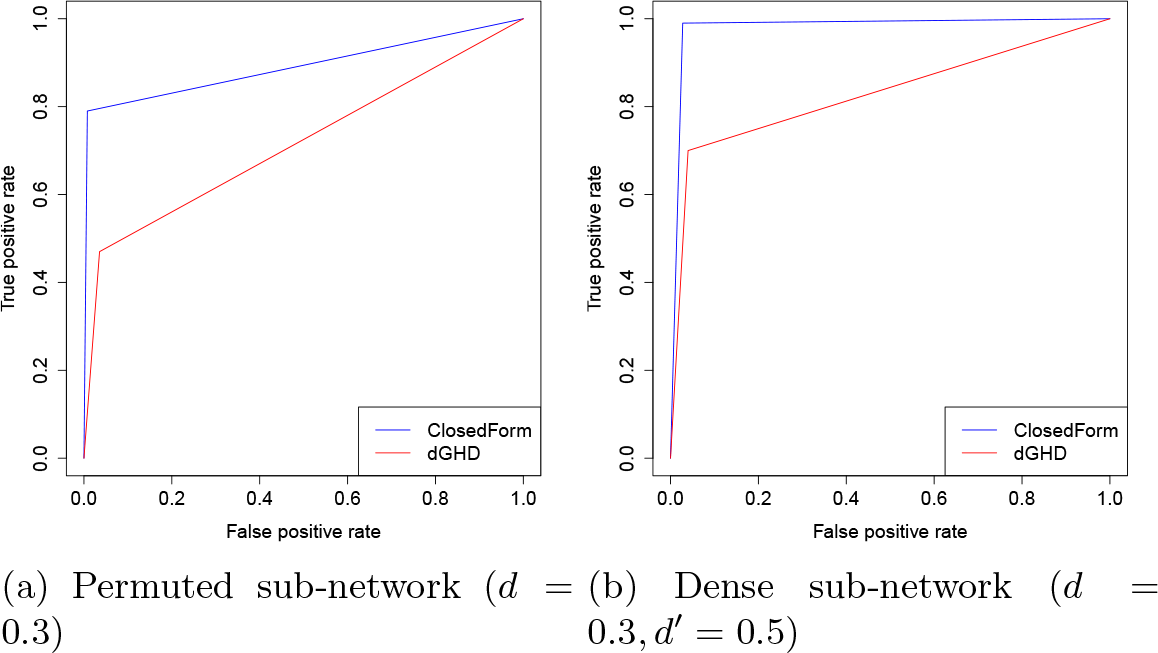
Comparison of proposed Closed-Form approach with dGHD algorithm. The plot of Closed-Form and dGHD methods are median plots w.r.t. to AUC metric out of 100 random runs. Clearly, the Closed-Form technique has better performance than dGHD algorithm.

We use the threshold 0.01 as cut-off for p-values in order to determine the true positives (TP) and true negatives (TN). We use median AUC value for the Closed-Form and dGHD techniques when comparing the ROC curves. We evaluate the true positive rate i.e.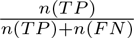 and the false positive rate i.e. 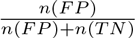 to estimate the ROC curve for these methods using the ‘pROC’ package in R. Here *n*(·) represents the total number of nodes. We also evaluated the area under the curve (AUC Mankiewicz (2004)) for the 100 runs of Closed-Form and dGHD methods.

**Closed-Form approach achieves better area under ROC curve in case of differential sub-networks formed by permuted nodes and subnetworks with higher density as shown in Figure 2.**

One of the reasons for relatively poor performance of the dGHD approach is that it has low true positive rate (TP) and a high false positive rate (FP) when the differential sub-network has more edges. This is also reflected by the relatively low Precision values for the dGHD algorithm in Table 2 when *d* = 0.3. From Figure 2b, we can observe that the median performance of both the dGHD and Closed-Form algorithm improves when the differential sub-network is denser than the remaining network.

**AUC value distributions for Closed-Form and dGHD techniques are statistically different**. For this experiment, we use the same set of networks as we used in the previous experiment and illustrate the results in (Supplementary Figure 2).

**Summary Table 2 highlights the computational efficiency and better predictive capabilities of the proposed techniques in comparison to dGHD algorithm**. For this comparison, we report the results obtained on 100 random runs of RG networks with *N* = 1000, *d* = 0.15 and *d* = 0.3 respectively, where the first 100 nodes are permuted. We also report results when the first 100 nodes form the denser differential sub-networks i.e. in experiments where *d* = 0.15 use *d*′ = 0.3 to form denser sub-network and where *d* = 0.3 use *d*′ = 0.5 to form denser sub-network. We also conducted experiments on undirected Power Law (PL) graphs using *N* = 1000 and *E* = 10, 000 with power law exponents *α* = {2,3} respectively. We permuted the first 100 nodes of each PL network (*B*) to form the permuted network (*A*). We performed 100 random runs and report the mean values for various evaluation metrics.

Table 2 compares the Closed-Form and dGHD techniques w.r.t. various standard evaluation metrics like AUC, Precision, Recall, Accuracy, Specificity, Kappa statistic and computational Time for all the simulation experiments. The evaluation metric Recall is equivalent to true positive rate used previously in our experiments. Higher values of these evaluation metrics represents better quality results. Here the time required by dGHD algorithm is normalized to 1 and the time required by the efficient implementation of the Closed-Form approach is scaled by the same normalization factor.

**Table 2:** Comparison of proposed Closed-Form approach with dGHD algorithm w.r.t. various evaluation metrics for random geometric (RG) and power law (PL) networks. Bold represents the best results.

| Parameters | Configuration | Method | AUC | Precision | Recall | Accuracy | Spec |
| --- | --- | --- | --- | --- | --- | --- | --- |
| | | | Mean ± Sd | Mean ± Sd | Mean ± Sd | Mean ± Sd | Mean |
| $d = 0.15$ | Permuted Subnet (RG) | **Closed-Form** | **0.935 ± 0.051** | **0.849 ± 0.037** | 0.846 ± 0.102 | **0.969 ± 0.011** | **0.983** |
| $d = 0.15$ | Permuted Subnet (RG) | **dGHD** | 0.926 ± 0.018 | 0.793 ± 0.021 | **0.878 ± 0.036** | 0.965 ± 0.005 | 0.974 |
| $d = 0.15, d' = 0.3$ | Denser Subnet (RG) | **Closed-Form** | **0.927 ± 0.048** | **0.839 ± 0.031** | 0.862 ± 0.098 | **0.969 ± 0.008** | **0.982** |
| $d = 0.15, d' = 0.3$ | Denser Subnet (RG) | **dGHD** | 0.922 ± 0.022 | 0.806 ± 0.027 | **0.868 ± 0.045** | 0.966 ± 0.006 | 0.977 |
| $d = 0.3$ | Permuted Subnet (RG) | **Closed-Form** | **0.877 ± 0.067** | **0.714 ± 0.075** | **0.789 ± 0.135** | **0.947 ± 0.016** | **0.975** |
| $d = 0.3$ | Permuted Subnet (RG) | **dGHD** | 0.724 ± 0.029 | 0.645 ± 0.049 | 0.577 ± 0.059 | 0.921 ± 0.007 | 0.971 |
| $d = 0.3, d' = 0.5$ | Denser Subnet (RG) | **Closed-Form** | **0.979 ± 0.005** | **0.771 ± 0.061** | **0.930 ± 0.082** | **0.965 ± 0.012** | **0.969** |
| $d = 0.3, d' = 0.5$ | Denser Subnet (RG) | **dGHD** | 0.848 ± 0.071 | 0.700 ± 0.038 | 0.731 ± 0.148 | 0.941 ± 0.010 | 0.964 |
| $\alpha = 2$ | Permuted Subnet (PL) | **Closed-Form** | **0.797 ± 0.046** | **0.307 ± 0.307** | 0.792 ± 0.099 | **0.801 ± 0.018** | **0.349** |
| $\alpha = 2$ | Permuted Subnet (PL) | **dGHD** | **0.797 ± 0.013** | 0.294 ± 0.009 | **0.809 ± 0.027** | 0.787 ± 0.008 | 0.333 |
| $\alpha = 3$ | Permuted Subnet (PL) | **Closed-Form** | **0.825 ± 0.019** | **0.345 ± 0.015** | **0.825 ± 0.035** | **0.826 ± 0.007** | **0.402** |
| $\alpha = 3$ | Permuted Subnet (PL) | **dGHD** | 0.808 ± 0.027 | 0.327 ± 0.018 | 0.799 ± 0.050 | 0.816 ± 0.008 | 0.375 |

We observe from Table 2 that the Closed-Form approach performs exceedingly well in case of experiments on denser RG networks (*d* = 0.3). For this configuration, in case of both permuted and denser differential sub-networks, the mean AUC of Closed-Form approach is at least 10% higher than the dGHD algorithm. This is also reflected in higher values of Precision (0.714 and 0.771) and Recall (0.789 and 0.930) metrics for Closed-Form approach in comparison to low values of Precision (0.645 and 0.7) and Recall (0.577 and 0.731) for the dGHD algorithm in case of these experiments. However, in case of sparser networks where its relatively easier to identify differential sub-networks Ruan *et al.*, (2015)), both the methods have similar predictive performance. Taken together these results show that the proposed Closed-Form approach outperforms dGHD technique w.r.t. various quality metrics like AUC, Precision, Recall, Specificity, Kappa and Time for both random geometric and power law graphs.

## 8 Case study

As a case study, we performed the differential sub-networks analysis of two gene regulatory networks re-constructed from the glioma dataset available on the TCGA Research Network (http://cancergenome.nih.gov). We recently reported that the integrative analysis of 1,122 glioma samples revealed the presence of seven groups with distinct molecular and clinical features (Ceccarelli *et al.*, 2016). In addition, we and others (Eckel-Passow*et al.*, 2015) showed that the majority of gliomas are divided into two main macro-categories according to the mutation of the gene IDH1. Therefore, our main biological question, that motivated the development of the reported methodology, was to identify the sub-networks of differentially activated transcription factors (TFs) in these two major conditions. We re-constructed two gene regulatory networks belonging to two different glioma subtypes: IDH-mutant and IDH-wild-type. Both networks were re-constructed with a four step procedure that extends ARACNE (Margolin *et al.*, 2006): i) Computation of mutual information between gene expression profiles to determine interaction between TFs and targets (Sales and Romualdi, 2011); ii) Data processing inequality to filter out indirect relationships (Margolin *et al.*, 2006), iii) Permutation test with 1, 000 re-samplings to keep only statistically significant relationships, and iv) Intersection with transcription factor binding sites to keep only relationships due to promoter binding.

We obtained two final networks consisting of 13, 683 unique connections for IDH-mutant and 14,158 for IDH-wild-type between 457 TFs and 4, 085 target genes. Using these networks, we construct the topological graphs as described in the Methods section for the 457 TFs. We then perform the proposed differential sub-network analysis to identify the TFs which are part of differential sub-networks in the topological graphs. Figure 3 shows the topmost differential sub-networks and Table 3 reports the topmost TFs which are part of differential sub-networks as detected by our algorithm. In the table, GHD and Mu (*μ*), represent the generalized hamming distance computed between networks without the transcription factor and the asymptotic mean *μ_π_* of GHD. The number of connections belonging in one network but not in the other is shown in the *Diff targets* column. It might happen that for some transcription factors such a difference is 0. This is because the networks under consideration are weighted and contribution of each node in the cGHD is dependent on the weighted degree of the node.

**Table 3:** The top most different transcription factors subnetworks detected between IDH-mutant and IDH-wild-type networks. The first four columns report differential measures in terms of p-value of the proposed differencing test, GHD computed between the two networks, the mean of the null GHD distribution, and the number of targets that belong exclusively in one network. The last three columns report the False Discovery Rate of the Fisher exact test obtained with a master regulator analysis, and the mean of transcription factor activity in IDH mut and wild-type. Transcription factor activity explains whether the transcription factor regulates directly (> 0) or inversely (< 0) its targets in the given condition.

| TF | Network differencing |  |  |  | Master Regulator Analysis |  |  |
| --- | --- | --- | --- | --- | --- | --- | --- |
|  | P-value | GHD | Mu | Diff targets | FDR | IDH mut activity | IDH WT activity |
| E2F1 | 0.951 | 0.059 | 0.059 | 35 | 1.000E+00 | -2.018 | -1.204 |
| ETV1 | 0.936 | 0.059 | 0.059 | 54 | 1.000E+00 | 1.670 | 1.161 |
| **RUNX3** | 0.837 | 0.060 | 0.060 | 55 | 7.192E-05 | -1.503 | -0.029 |
| CREB1 | 0.825 | 0.059 | 0.059 | 42 | 1.000E+00 | 1.097 | 0.924 |
| **FOXD2** | 0.740 | 1.000 | 0.300 | 71 | 8.538E-09 | -3.680 | -1.274 |
| **FOXJ2** | 0.740 | 1.000 | 0.300 | 119 | 7.424E-03 | 0.850 | -0.537 |
| **MEIS1** | 0.740 | 1.000 | 0.300 | 82 | 7.475E-11 | -1.181 | 1.064 |
| MTF1 | 0.740 | 1.000 | 0.300 | 107 | 1.000E+00 | 0.401 | 0.389 |
| **KLF13** | 0.735 | 1.000 | 0.278 | 80 | 8.086E-04 | 1.014 | -0.255 |
| **SOX10** | 0.726 | 0.060 | 0.060 | 40 | 1.573E-07 | 0.858 | -1.130 |
| **STAT3** | 0.717 | 0.059 | 0.059 | 41 | 1.112E-31 | -0.318 | 1.335 |
| **IRF3** | 0.685 | 0.840 | 0.455 | 91 | 2.356E-13 | -1.505 | 0.142 |
| **HOXD13** | 0.618 | 0.060 | 0.060 | 44 | 8.705E-07 | -1.840 | -0.223 |
| ZNF354C | 0.616 | 0.059 | 0.059 | 39 | 1.000E+00 |  |  |
| **ZIC1** | 0.534 | 0.059 | 0.059 | 0 | 3.319E-19 | -2.752 | 0.475 |
| **HOXA2** | 0.513 | 0.060 | 0.060 | 24 | 2.541E-02 | -1.388 | 0.201 |
| **FOXO1** | 0.459 | 0.060 | 0.059 | 2 | 2.572E-02 | -2.344 | -0.687 |
| DLX6 | 0.416 | 0.060 | 0.060 | 23 | 1.000E+00 |  |  |
| MAFG | 0.414 | 0.862 | 0.467 | 60 | 1.000E+00 | 0.739 | -0.100 |
| NR4A2 | 0.394 | 0.060 | 0.060 | 6 | 1.000E+00 | -0.169 | -0.318 |
| PAX6 | 0.331 | 0.060 | 0.059 | 39 | 9.057E-01 | 2.209 | 1.416 |
| MEF2D | 0.326 | 0.060 | 0.060 | 56 | 1.567E-01 | 0.406 | -0.583 |
| NR1H2 | 0.271 | 0.059 | 0.059 | 44 | 1.000E+00 | -2.363 | -0.399 |
| RFX1 | 0.259 | 0.060 | 0.061 | 26 | 1.768E-01 | -0.060 | 0.958 |
| STAT4 | 0.255 | 0.848 | 0.486 | 78 | 9.025E-01 | -0.929 | -1.049 |
| **SIX4** | 0.226 | 0.060 | 0.059 | 7 | 5.592E-03 | 2.040 | 0.004 |
| GLIS2 | 0.196 | 0.060 | 0.061 | 24 | 4.905E-01 | 0.332 | -0.699 |
| **OTP** | 0.190 | 0.060 | 0.060 | 9 | 2.156E-05 | -1.017 | 0.911 |
| **HOXB4** | 0.159 | 0.060 | 0.060 | 17 | 6.416E-03 | -2.019 | -0.345 |
| BACH1 | 0.145 | 0.060 | 0.060 | 57 | 1.000E+00 | -0.565 | 0.223 |
| **MBD2** | 0.139 | 0.820 | 0.495 | 76 | 2.330E-10 | -1.488 | 0.070 |
| IRF9 | 0.132 | 0.060 | 0.060 | 15 | 1.000E+00 | 0.302 | 0.675 |
| NR2C1 | 0.110 | 0.061 | 0.060 | 19 | 1.000E+00 | -0.147 | -0.380 |
| **KLF6** | 0.107 | 0.060 | 0.061 | 23 | 9.536E-07 | -1.378 | 0.333 |
| HMBOX1 | 0.094 | 0.196 | 0.191 | 28 | 9.184E-01 | 0.367 | -0.545 |
| CREM | 0.094 | 0.765 | 0.506 | 51 | 1.408E-01 | -1.594 | -0.324 |
| HOXA9 | 0.092 | 0.061 | 0.060 | 21 | 2.526E-01 | -2.820 | -1.082 |
| **ZIC2** | 0.086 | 0.198 | 0.192 | 39 | 3.633E-05 | -0.786 | 0.468 |
| **MEOX2** | 0.078 | 0.200 | 0.194 | 53 | 2.960E-29 | -1.420 | 0.749 |
| ETV4 | 0.075 | 0.060 | 0.061 | 17 | 1.661E-01 | -0.874 | 0.782 |
| HOXA13 | 0.071 | 0.061 | 0.060 | 61 | 1.000E+00 | -2.053 | -1.481 |
| **SMAD1** | 0.069 | 0.202 | 0.195 | 51 | 1.450E-04 | -0.013 | 1.493 |
| RFX4 | 0.060 | 0.204 | 0.197 | 28 | 1.000E+00 | -0.278 | 0.819 |
| **ZBTB12** | 0.055 | 0.206 | 0.199 | 37 | 4.556E-03 | 1.304 | -0.362 |
| **STAT1** | 0.051 | 0.060 | 0.061 | 18 | 7.839E-29 | -1.002 | 1.002 |

To asses the biological validity, we also assembled a global glioma network using all the available transcriptional profiles using the same method described above and performed a master regulator analysis (Carro *et al.*, 2010) with respect to the molecular phenotype under investigation, *i.e.* genes differential expressed between IHD mutant and wild type. Master regulator analysis is extensively adopted to identify TFs that act as principal regulators in driving the phenotype from one condition to another. The last three columns of the table show the master regulator analysis results for each TF (in boldface the most significant master regulators).

Interestingly, among the topmost TFs (out of 457) forming the differential sub-networks, we found several genes known to have a central role in controlling specific glioma subtypes as well as novel candidates that deserve further biological validation. In particular, our proposed algorithms reveals that the subnetwork of STAT3 is one the most different between IDH-mutant and IDH-wild-type networks and a particularly significant Master Regulator of this wild-type phenotype. Members of our group have previously shown that STAT3, together with C/EBP*β*, is a key regulator of the mesenchymal differentiation and predicts the poor clinical outcome of IDH-wild-type gliomas (Carro *et al.*, 2010). Another key regulator of the IDH-wild-type gliomas was recently reported by using an integrative functional copy number analysis is the set of HOXA genes (Ceccarelli *et al.*, 2016). Moreover, another key network hub that our algorithm detects as different is SOX10 which appears to be an active master regulator of the IDH-mutant phenotype. We recently reported that the GCIMP-low subgroup in the IDH-mutant cohort can mediated by loss of CpG methylation and binding of SOX factors (Ceccarelli *et al.*, 2016). Furthermore, our algorithm identifies methyl-CpG-binding domain protein 2 (MBD2) as a main differential network hub. In particular, MBD2 has no links in the IDH-wild-type network whereas it is highly connected in the IDH-mutant network which is characterized by the CpG island methylator phenotype (GCIMP) (Noushmehr *et al.*, 2010). MBD2 is a mediator of the epigenetic gene regulation and its role in glioblastoma is being studied as its over-expression may drive tumor growth by suppressing the anti-angiogenic activity of key tumor suppressors (Zhu *et al.*, 2011).

## 9 Conclusions

The comparison of gene expression profiles across different phenotypes is enabling the discovery of novel biomarkers for prognosis or diagnosis. They hold the key to identify novel targets for therapeutical intervention. In this paper we proposed an improvement to the state-of-the-art for comparing two labeled graphs that are representative of two conditions (e.g. the macro-categories according to the mutation of the gene IDH1 in our case study) and identifying statistically significant differences in their topology. We used the centralized GHD (cGHD) metric (Ruan *et al.*, 2015) to calculate the distance between the two labelled networks. We proposed a Closed-Form approach, an improvement to the dGHD algorithm, to detect localized topological differences between paired networks. The Closed-Form approach calculates the closed-form contri-bution of each node in the cGHD metric and efficiently removes nodes with the smaller contributions in the cGHD value. From our experiments on scale free random geometric networks, we discovered that the Closed-Form approach was 10-15x faster than Original method from a computational complexity point of view. For differential sub-network analysis in very sparse paired graphs, both the Closed-Form and Original methods had good predictive performance. They reached mean AUC values of ≈ 0.932 and ≈ 0.924 respectively for 100 random runs of simulation experiments. However, for relatively denser networks, the Closed-Form approach outperformed the Original method. The proposed method achieved a mean AUC of ≈ 0.915 while the Original technique reached a mean AUC of ≈ 0.79. The Closed-Form approach also achieved much higher Precision, Recall and Kappa values in comparison to the Original method for relatively denser networks. We applied our algorithm to detect the main differences between the networks of IDH-mutant and IDH-wild-type glioma tumors and show that it correctly selects sub-networks centered on important key regulators of these two different subtypes. In addition its application highlights novel candidates, such as MBD2, that can be the subject of further biological validations.

**Figure 3:**
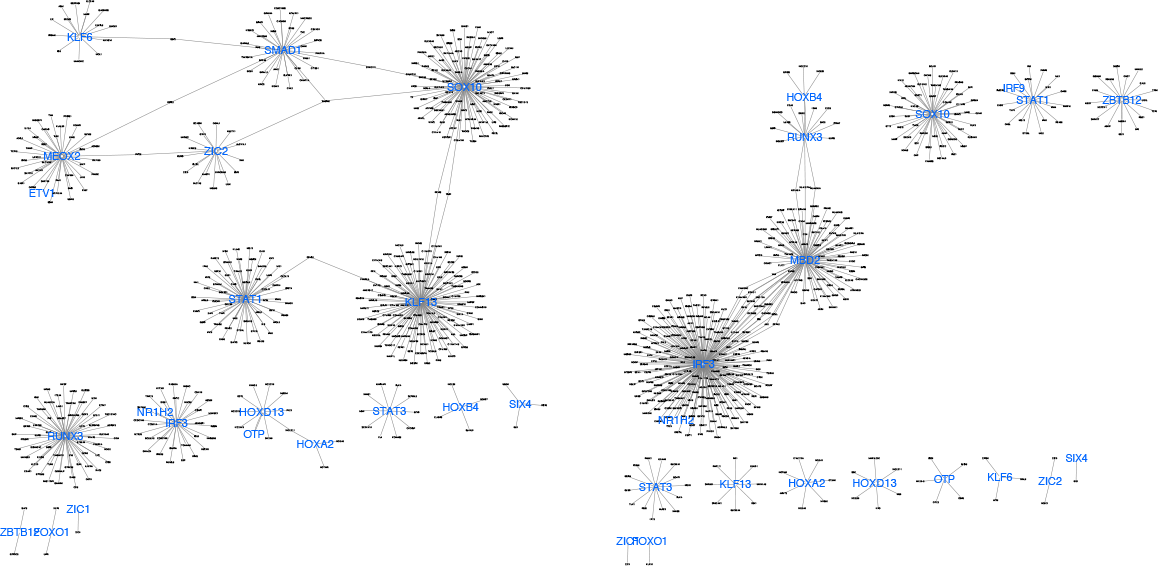
Differential sub-networks in IDH mutant (rigth) and IDH wild type (left). In blue the most different transcription factors.

